# Hyperexcitability and loss of feedforward inhibition in the *Fmr1*KO lateral amygdala

**DOI:** 10.1101/2020.04.21.053652

**Authors:** E. Mae Guthman, Matthew N. Svalina, Christian A. Cea-Del Rio, J. Keenan Kushner, Serapio M. Baca, Diego Restrepo, Molly M. Huntsman

## Abstract

Fragile X Syndrome (FXS) is a neurodevelopmental disorder characterized by intellectual disability, autism spectrum disorders (ASDs), and anxiety disorders. The disruption in the function of the *FMR1* gene results in a range of alterations in cellular and synaptic function. Previous studies have identified dynamic alterations in inhibitory neurotransmission in early postnatal development in the amygdala of the mouse model of FXS. Yet little is known how these changes alter microcircuit development and plasticity in the lateral amygdala (LA). Using whole-cell patch clamp electrophysiology, we demonstrate that principal neurons (PNs) in the LA exhibit hyperexcitability with a concomitant increase in the synaptic strength of excitatory synapses in the BLA. Further, reduced feed-forward inhibition appears to enhance synaptic plasticity in the FXS amygdala. These results demonstrate that plasticity is enhanced in the amygdala of the juvenile *Fmr1* KO mouse and that E/I imbalance may underpin anxiety disorders commonly seen in FXS and ASDs.

## INTRODUCTION

Fragile X Syndrome (FXS) is the most common monogenic form of intellectual disability. FXS is a neurodevelopmental disorder (NDD) broadly characterized by neurologic and psychiatric disorders such as attention deficit hyperactivity disorder, anxiety, social avoidance, increased incidence of seizures and epilepsy, and autism spectrum disorders (ASDs)(Hagerman *et al.*, 2009). In humans, FXS is caused by a repeat expansion mutation in the *FMR1* gene that encodes fragile x mental retardation protein (FMRP)(Liu *et al.*, 2018). Trinucleotide repeat expansion results in hypermethylation at the *FMR1* locus and subsequent transcriptional silencing of FMRP (Fu *et al.*, 1991). FMRP is an RNA binding protein with a known role in regulating messenger RNA translation during synaptic development (Chen *et al.*, 2003; Darnell *et al.*, 2011). The dysregulation of protein synthesis observed in the pathogenesis of FXS is known to result in significant defects in neuronal development, synaptic and circuit function (Contractor *et al.*, 2015).

In FXS, profound alterations in excitatory and inhibitory neurotransmission have been found across multiple brain regions including somatosensory cortex and the basolateral amygdala (BLA) (Huber *et al.*, 2002; Olmos-Serrano *et al.*, 2010; Contractor *et al.*, 2015). Indeed, accumulating evidence directly implicates BLA dysfunction as a key component of many behavioral manifestations in FXS as well as NDDs (Baron-Cohen *et al.*, 2000; Hessl *et al.*, 2004; Bauman & Kemper, 2005; Dalton *et al.*, 2005). Amygdala-based behaviors, including anxiety disorders and social withdrawal, are commonly diagnosed psychiatric disorders in individuals with FXS and ASDs (Tsiouris & Brown, 2004; Turk *et al.*, 2005; Cordeiro *et al.*, 2011). In NDDs such as FXS, patients have increased anxiety and an increased retention of fearful memories (Turk *et al.*, 2005). The adherence to fearful memories dictates the emotional state of the patient (Turk *et al.*, 2005) and likely exacerbates already increased anxiety levels (Meredith *et al.*, 2012). Further, patients with intellectual disabilities can exhibit stress and anxiety from an overactive response to fearful memories, similar to post traumatic stress and panic disorders (Turk *et al.*, 2005; Roberts *et al.*, 2009).

The amygdala is a grouping of many distinct, heterogeneous nuclei responsible for the integration and processing of information with emotional and social salience (Duvarci & Pare, 2014; Janak & Tye, 2015; Li *et al.*, 2017). Specifically, a large body of work has identified the BLA as the main site of synaptic plasticity underlying the acquisition, expression, and extinction of sensory-threat associations with the BLA also implicated in neuropsychiatric diseases such as anxiety disorders (Duvarci & Pare, 2014; Janak & Tye, 2015). At the cellular level, the BLA is composed of excitatory principal neurons (PNs) and a diverse population of GABAergic inhibitory interneurons (INs) (McDonald, 1984; Sah *et al.*, 2003; Duvarci & Pare, 2014). These local PNs undergo input-specific, activity-dependent plastic changes in response to co-occurring threatening and sensory stimuli (Quirk *et al.*, 1995; McKernan & Shinnick-Gallagher, 1997; Schoenbaum *et al.*, 1999; Nabavi *et al.*, 2014; Grewe *et al.*, 2017; Kim & Cho, 2017; Kasugai *et al.*, 2019). Importantly, these co-occurring stimuli depolarize and drive ensembles of BLA PNs to fire action potentials, and this excitation is necessary for learning to occur (Rosenkranz & Grace, 2002; Wolff *et al.*, 2014; Grewe *et al.*, 2017). The local circuit INs play an important role regulating the synaptic plasticity underlying this *in vivo* learning process (Bissière *et al.*, 2003; Wolff *et al.*, 2014). Our group and others have shown this inhibitory control is exerted via a feedforward circuit motif from somatostatin-expressing (Sst) INs which modulates long-term potentiation (LTP) at cortical and thalamic afferent synapses onto BLA PNs (Smith *et al.*, 2000; Bissière *et al.*, 2003; Tully *et al.*, 2007; Wolff *et al.*, 2014; Unal *et al.*, 2014; Bazelot *et al.*, 2015; Ito *et al.*, 2019; Guthman *et al.*, 2020). Thus, feedforward inhibition-gated LTP is an underlying circuit mechanism for the acquisition of threat conditioning.

In FXS, profound alterations in the GABAergic system have been previously identified in cortex, hippocampus, brainstem, and BLA (El Idrissi *et al.*, 2005; D’Hulst *et al.*, 2006; Gibson *et al.*, 2008; Olmos-Serrano *et al.*, 2010; Paluszkiewicz *et al.*, 2011; Vislay *et al.*, 2013; Martin *et al.*, 2014). Specifically, prior work from our group has shown reductions in both tonic and phasic inhibitory neurotransmission as well as GABA availability have been observed in the rodent lateral amygdala (LA) (Olmos-Serrano *et al.*, 2010; Martin *et al.*, 2014), the BLA subnucleus where these fundamental plasticity processes first occur in the canonical BLA circuit (Duvarci & Pare, 2014; Janak & Tye, 2015).

In the present study, we combined whole-cell patch clamp electrophysiology to explore the intrinsic properties of LA PNs, local microcircuit excitation-inhibition (E/I) balance, and synaptic plasticity. We found that excitatory PNs in the *Fmr1*KO LA show marked hyperexcitability compared to wildtype (WT) animals. Consistent with the role of feedforward inhibition (FFI) in gating LTP in the LA, we show a correlated loss of FFI and enhanced LTP in *Fmr1*KO LA. These results demonstrate that altered E/I balance in mice enhances synaptic plasticity in LA and may underpin behavioral disorders seen in both children with FXS and ASDs.

## RESULTS

### LA PNs in *Fmr1*KO mice exhibit marked hyperexcitability

To examine potential differences in neuronal excitability, we prepared acute coronal brain slices containing the BLA. We performed whole cell patch-clamp recordings of LA PNs and compared their intrinsic biophysical properties across wildtype (WT) and *Fmr1*KO juvenile mice (postnatal days 21-35). In these experiments, we measured 18 membrane properties (Fig. 1, Table 1) by examining voltage responses to both a ramped and rectangle current injections (See Materials & Methods).

**Table 1.** Differences in active and passive membrane properties among LA PNs in WT and Fmr1KO mice. Normal data are presented as mean ± s.e.m. with differences tested using an unpaired t-test. Non-normal data are presented as median and IQR with differences tested using a Mann-Whitney U (MWU) test.

|  | WT PN<sub>s</sub> (n = 22) |  | *Fmr1*KO PN<sub>s</sub> (n = 24) |  | Statistical Comparisons |
| --- | --- | --- | --- | --- | --- |
|  | *Mean/*<br>*Median* | *Variance* | *Mean/*<br>*Median* | *Variance* |  |
| **Membrane Resistance (MW)** | 159.89 | 132.75/211.01 | 218.94 | 190.68/347.18 | $p = 0.0016$, MWU test |
| **$\tau_{\text{Membrane}}$ (ms)** | 25.24 | 23.08/30.38 | 30.92 | 27.50/34.49 | $p = 0.0016$, MWU test |
| **Max. Firing Rate (Hz)** | 25.66 | $\pm 1.99$ | 31.37 | $\pm 1.25$ | $p = 0.018$, unpaired t-test |
| **Action Potential Halfwidth (ms)** | 0.95 | 0.90/1.15 | 1.18 | 1.00/1.63 | $p = 0.0034$, MWU test |
| **Rheobase Current (pA)** | 46.83 | $\pm 3.26$ | 50.97 | $\pm 2.44$ | $p = 0.0013$, unpaired t-test |
| **Action Potential Threshold (mV)** | -35.75 | $\pm 0.69$ | -35.50 | $\pm 0.91$ | $p = 0.83$, unpaired t-test |
| **Latency to First Action Potential (ms)** | 125.60 | 111.30/167.40 | 137.15 | 110.40/175.30 | $p = 0.53$, MWU test |
| **Firing Rate Adaptation** | 0.54 | $\pm 0.031$ | 0.54 | $\pm 0.024$ | $p = 0.93$, unpaired t-test |
| Action Potential Broadening | 1.12 | 1.08/1.25 | 1.20 | 1.11/1.34 | $p = 0.16$,<br>MWU test |
| Action Potential Amplitude (mV) | 69.56 | $\pm 1.79$ | 66.17 | $\pm 2.68$ | $p = 0.31$,<br>unpaired t-test |
| Amplitude Adaptation | 0.96 | 0.91/0.98 | 0.96 | 0.93/0.99 | $p = 0.77$,<br>MWU test |
| AHP magnitude (mV) | 17.57 | $\pm 0.63$ | 18.65 | $\pm 0.59$ | $p = 0.22$,<br>unpaired t-test |
| DAHP (mV) | -2.76 | -2.44/-3.99 | -3.61 | -2.38/-3.61 | $p = 0.52$,<br>MWU test |
| AHP Latency (ms) | 46.83 | $\pm 3.26$ | 50.97 | $\pm 2.44$ | $p = 0.31$,<br>unpaired t-test |
| Hyperpolarization-induced Sag (%) | 6.24 | 5.19/7.96 | 8.32 | 4.31/9.59 | $p = 0.62$,<br>MWU test |
| Rebound Action Potentials | 0.00 | 0.00/0.00 | 0.00 | 0.00/0.00 | $p = 1.00$,<br>MWU test |

**Figure 1.**
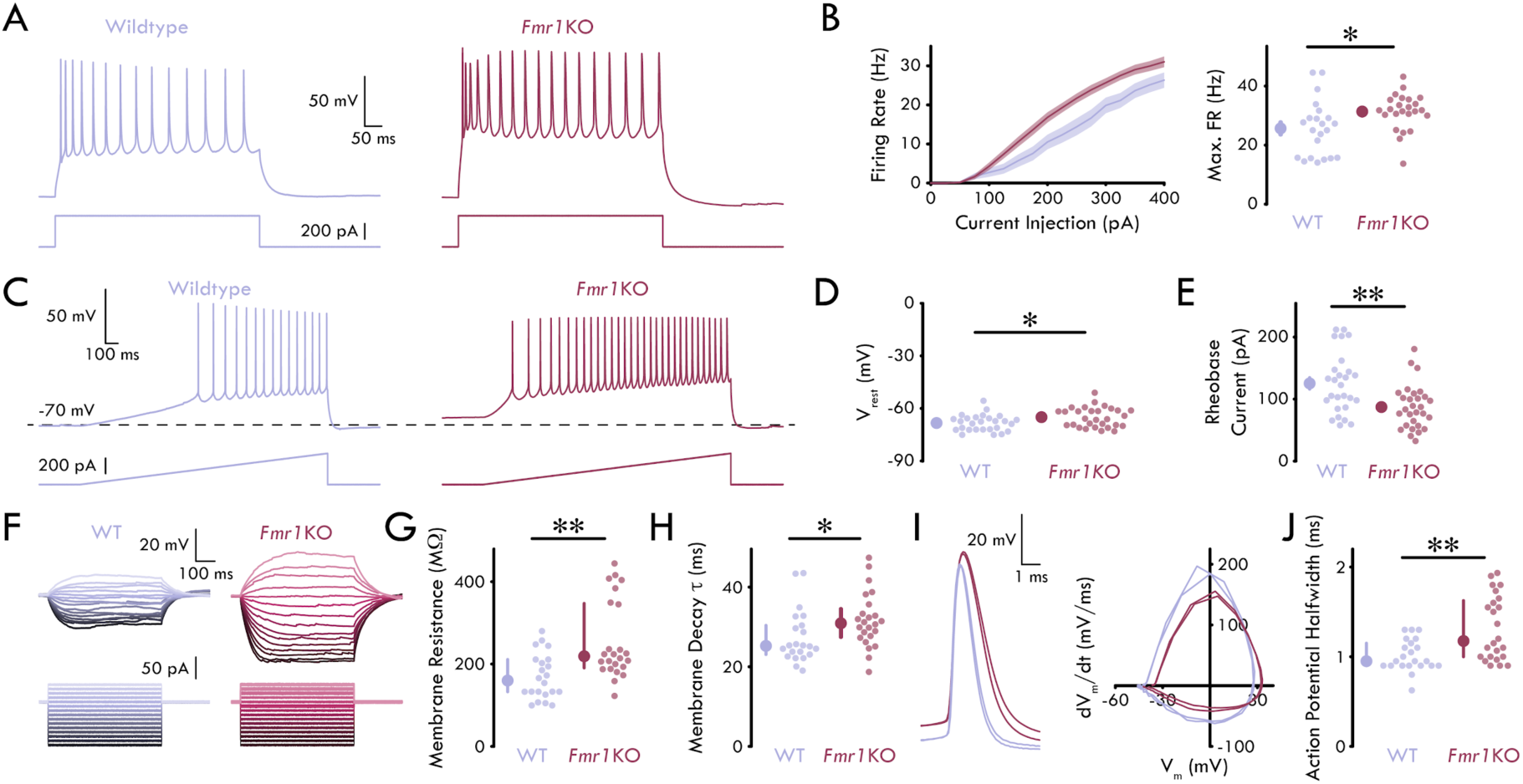
Hyperexcitability of PNs in *Fmr1*KO LA. **(A)** Representative traces of maximum firing rate response to rectangular current injections in WT and *Fmr1*KO LA PNs **(B)** Left: Mean firing rate of LA PNs. Shading shows s.e.m. Right: Maximum firing rate of LA PNs is greater in *Fmr1*KO compared to WT LA (unpaired t-test: *p* = 0.018; n_WT_ = 22 neurons, 3 mice; n_*Fmr1*KO_ = 24, 4) **(C)** Representative traces of voltage response to a ramped current injection in WT and *Fmr1*KO LA PNs **(D)** *Fmr1*KO LA PNs have a more depolarized V_rest_ compared to WT LA PNs (unpaired t-test: p = 0.019; n_WT_ = 27, 4; n_Fmr1KO_ = 29, 4) **(E)** *Fmr1*KO LA PNs have a lower rheobase current from rest compared to WT LA PNs (unpaired t-test: *p* = 0.0013; n_WT_ = 27, 4; n_*Fmr1*KO_ = 29, 4) **(F)** Representative traces of voltage responses to intermediate current injection traces used to determine membrane resistance and decay τ (−100 to +100 pA; Δ10 pA) **(G)** Membrane resistance is increased in *Fmr1*KO LA PNs compared to WT LA PNs (Mann-Whitney U test: *p* = 0.0016; n_WT_ = 22, 3; n_*Fmr1*KO_ = 24, 4) **(H)** Membrane decay τ is increased in *Fmr1*KO LA PNs compared to WT LA PNs (Mann-Whitney U test: *p* = 0.0016; n_WT_ = 22, 3; n_*Fmr1*KO_ = 24, 4) **(I)** Left: Representative action potential traces from a WT and *Fmr1*KO LA PN at rheobase current injection. Right: Phase plot for the same action potentials **(J)** *Fmr1*KO LA PNs have broader action potential halfwidths compared to WT LA PNs (Mann-Whitney U test: *p* = 0.0034; n_WT_ = 22, 3; n_*Fmr1*KO_ = 24, 4) Summary statistics in B, D, and E presented as mean ± s.e.m. Summary statistics in H and J presented as median with IQR. *p < 0.05, **p < 0.01.

We observed significant differences in both active and passive membrane properties of LA PNs in *Fmr1*KO compared to WT animals. Specifically, depolarizing current injections drove increased action potential firing rates in LA PNs of *Fmr1*KO compared to WT animals (Fig. 1A, B; unpaired t-test: *p* = 0.018). Additionally, PNs in the LA of *Fmr1*KO mice exhibited a higher resting membrane potential (V_rest_) and a lower rheobase current compared to PNs in WT mice (Fig. 1C-E; V_rest_, unpaired t-test: *p* = 0.019; rheobase current: unpaired t-test: *p* = 0.0013). Further, LA PNs in *Fmr1*KO mice showed increased membrane resistance (R_m_), increased membrane decay τ, and action potential halfwidth (Fig. 1 F-J; R_m_, Mann-Whitney U [MWU] test: *p* = 0.0016; decay τ, MWU test: *p* = 0.0016; halfwidth, MWU test: *p* = 0.0034). No other membrane property comparisons reached statistical significance (Table 1). Overall, these data reveal increased intrinsic membrane excitability in the LA PNs of *Fmr1*KO compared to WT mice.

### Alterations in spontaneous excitation and inhibition in *Fmr1*KO LA

Previous studies from our group identified defects in BLA inhibitory neurotransmission such that the frequency and amplitude of both phasic and tonic inhibitory postsynaptic currents (IPSCs) are reduced during the P21-35 development timepoint (Olmos-Serrano *et al.*, 2010; Vislay *et al.*, 2013; Martin *et al.*, 2014). One possible explanation for this reduction could be that it is a homeostatic response to a concomitant change in spontaneous excitatory postsynaptic currents (sEPSCs). However, our previous studies on synaptic transmission were done in the presence of NMDA and AMPA receptor antagonists (D-APV and DNQX, respectively) to induce a complete excitatory blockade and isolate spontaneous IPSCs (sIPSCs) (Olmos-Serrano *et al.*, 2010; Vislay *et al.*, 2013). To study how loss of *Fmr1* contributes to both spontaneous glutamatergic and GABAergic synaptic transmission in intact LA, we performed patch-clamp recordings in local PNs of *Fmr1*KO and WT animals using a cesium-methanesulfonate based intracellular solution. This solution allows us to voltage clamp the PNs at -70 mV to isolate sEPSCs and 0 mV to isolate sIPSCs (Fig. 2A, F).

**Figure 2.**
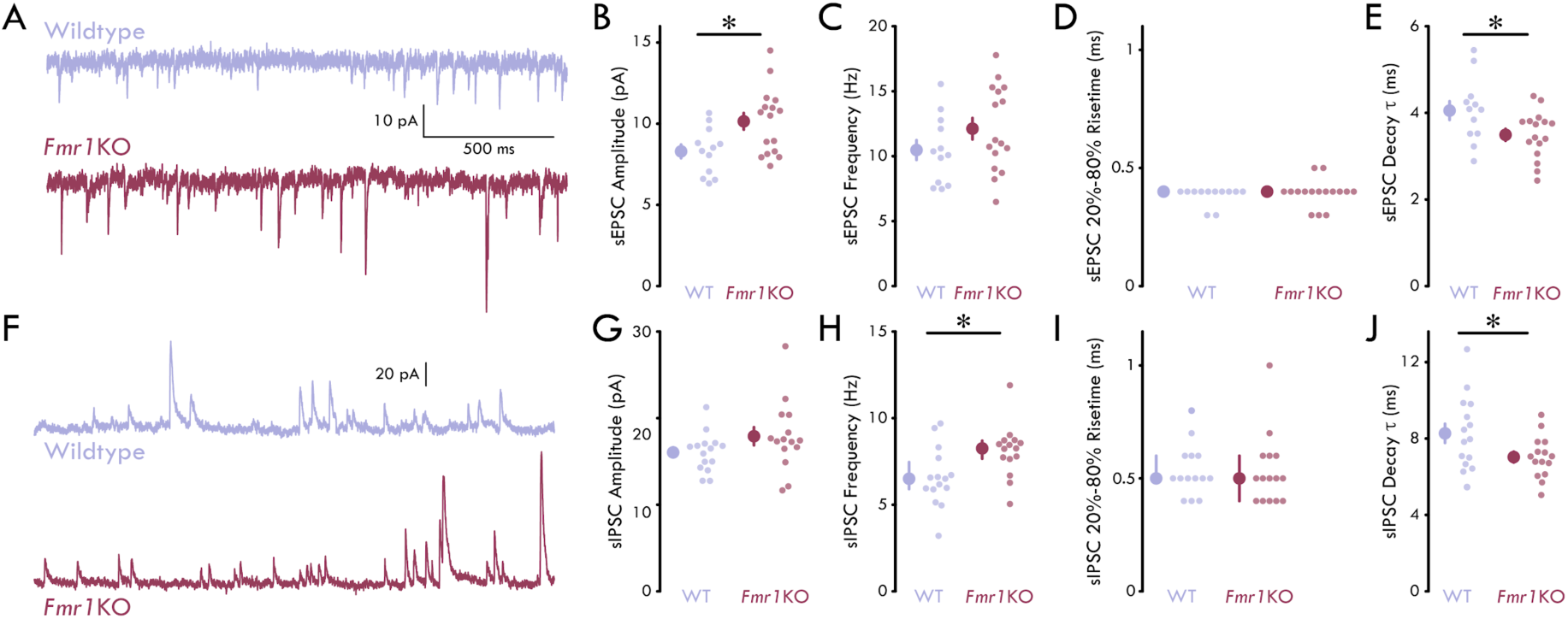
Impaired spontaneous excitatory and inhibitory postsynaptic currents in *Fmr1*KO LA. **(A)** Representative current traces from LA PNs held at -70 mV **(B)** sEPSC amplitude is increased in LA PNs of *Fmr1*KO mice (unpaired t-test: *p* = 0.013; n_WT_ = 12 neurons, 7 mice; n_*Fmr1*KO_ = 16, 5) **(C)** No significant difference in sEPSC frequency in LA PNs between WT and *Fmr1*KO mice (unpaired t-test: *p* = 0.17; n_WT_ = 12, 7; n_*Fmr1*KO_ = 16, 5) **(D)** No significant difference in sEPSC risetime in LA PNs between WT and *Fmr1*KO mice (MWU test: *p* = 0.65; n_WT_ = 12, 7; n_*Fmr1*KO_ = 16, 5) **(E)** sEPSC decay τ is reduced in LA PNs of *Fmr1*KO mice (unpaired t-test: *p* = 0.031; n_WT_ = 12, 7; n_*Fmr1*KO_ = 16, 5) **(F)** Representative current traces from LA PNs held at 0 mV **(G)** No significant difference in sIPSC amplitude in LA PNs between WT and *Fmr1*KO mice (unpaired t-test: *p* = 0.13; n_WT_ = 15, 7; n_*Fmr1*KO_ = 15, 4) **(H)** sIPSC frequency is increased in LA PNs of *Fmr1*KO mice (MWU test: *p* = 0.016; n_WT_ = 15, 7; n_*Fmr1*KO_ = 15, 4) **(I)** No significant difference in sIPSC risetime in LA PNs between WT and *Fmr1*KO mice (MWU test: *p* = 0.50; n_WT_ = 15, 7; n_*Fmr1*KO_ = 15, 4) **(J)** sIPSC decay τ is reduced in LA PNs of *Fmr1*KO mice (unpaired t-test: *p* = 0.039; n_WT_ = 15, 7; n_*Fmr1*KO_ = 15, 4) Summary statistics in C, E, G, and J presented as mean ± s.e.m. Summary statistics in D, H, and I presented as median with IQR. *p < 0.05.

We found that sEPSCs from PNs in *Fmr1*KO LA showed an increase in sEPSC amplitude relative to WT controls (Fig. 2B; unpaired t-test, *p* = 0.013). We found no differences in the frequency or 20%-80% risetime (Fig. 2C, D; frequency: unpaired t-test, *p* = 0.17; risetime: MWU test, *p* = 0.65). Additionally, sEPSCs from PNs in *Fmr1*KO showed a significantly decreased decay τ (Fig. 2E; unpaired t-test, *p* = 0.031). When we examined sIPSCs, we found no change in sIPSC amplitude; however, we found an increase in the frequency of sIPSCs onto local PNs (Fig. 2G, H; amplitude: unpaired t-test, *p* = 0.13; frequency: MWU test, *p* = 0.016). As with sEPSCs, we found no significant difference in the 20%-80% risetime of PNs in *Fmr1*KO and WT LA (Fig. 2I; MWU test, *p* = 0.50). However, LA PNs in *Fmr1*KO mice exhibited a decrease in sIPSC decay (Fig. 2J; unpaired t-test, *p* = 0.039). Taken together, these data identify differences in pre- and postsynaptic modulation of excitatory and inhibitory neurotransmission in the LA of *Fmr1*KO mice.

### FFI is reduced in *Fmr1*KO LA

Disrupted E/I balance of neuronal networks is a hallmark of NDDs (Nelson & Valakh, 2015). In FXS, this manifests as an increased prevalence of anxiety, epilepsy, and attention deficit and hyperactivity (Musumeci *et al.*, 1999; Rogers *et al.*, 2001; Clifford *et al.*, 2007). To further test the hypothesis that loss of *Fmr1* leads to a disruption in local circuit E/I balance, we measured the amplitudes of evoked EPSCs and IPSCs in LA PNs following stimulation of the internal capsule. The internal capsule carries thalamic afferents to the LA (LeDoux *et al.*, 1991). We focused on this afferent synapse as it is a major site of the input-specific LTP that underlies the acquisition of classical Pavlovian threat conditioning *in vivo* (McKernan & Shinnick-Gallagher, 1997; Namburi *et al.*, 2015).

Similar to the spontaneous synaptic event experiments above, we isolated evoked EPSC and IPSCs by voltage clamping LA PNs at -70 and 0 mV, respectively. To determine how loss of *Fmr1* affects feedforward excitatory and inhibitory drive onto LA PNs, we recorded evoked EPSCs and IPSCs over a stimulus intensity range of 0 - 100 µA (Figure 3A, B). We found that loss of *Fmr1* had no effect on feedforward excitatory drive to LA PNs as measured by either the stimulation intensity for the half-maximal EPSC amplitude or the slope of the input-output function (Fig. 3A-D; halfmax stimulation intensity: unpaired t-test, *p* = 0.080; input-output slope: unpaired t-test, *p* = 0.94). However, we found that feedforward inhibitory drive was reduced in LA PNs of *Fmr1*KO mice. Specifically, we found that loss of *Fmr1* led to an increase in the stimulation intensity for half-maximal IPSC amplitude in LA PNs (Fig. 3G; unpaired t-test, *p* = 0.017). There was no effect of loss of *Fmr1* on the slope of the input-output function (Fig. 3H; unpaired t-test, *p* = 0.62). These data indicate that a greater amount of activation of the thalamic afferents to LA is needed to drive similar FFI onto PNs in *Fmr1*KO compared to WT mice. However, the lack of change in input-output slope indicates that once afferent activity is sufficient to elicit IPSCs in the postsynaptic PNs, the IPSC amplitudes increase as a similar function of afferent activity. This finding is in accordance with prior work showing increased feedforward E/I balance in cortical microcircuits *Fmr1*KO mice at the same developmental time point (Antoine *et al.*, 2019).

**Figure 3.**
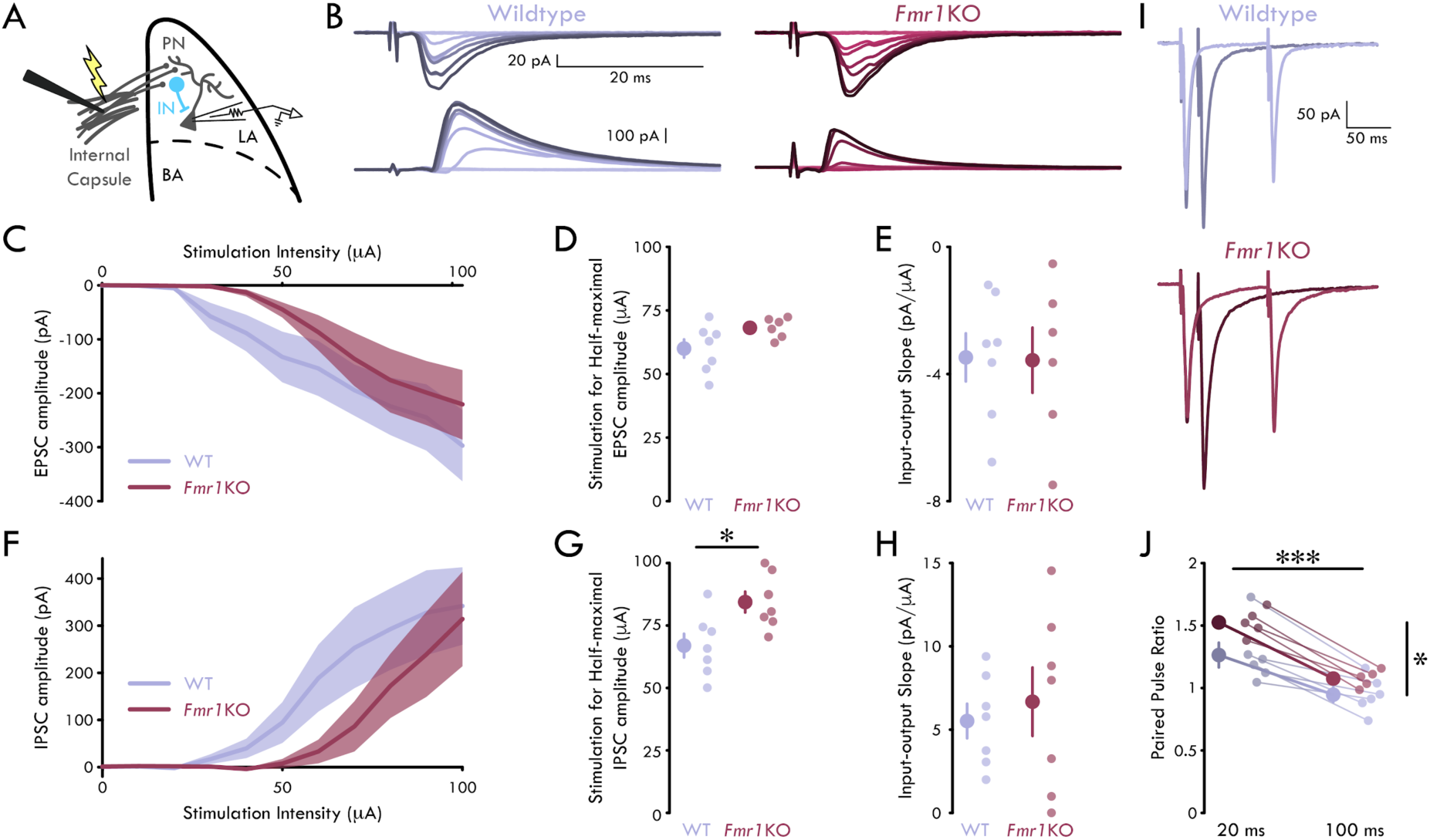
Reduced feedforward inhibition in *Fmr1*KO LA. **(A)** Experimental schematic. **(B)** Representative mean traces of EPSCs (top) and IPSCs (bottom) in LA PNs following internal capsule stimulation. Color scales with stimulation intensity (light to dark: 0 - 100 µA) **(C)** Mean evoked EPSC amplitude as a function of stimulation intensity. Shading shows s.e.m. **(D)** Stimulation for half-maximal EPSC amplitude is not significantly different for LA PNs from WT and *Fmr1*KO mice (unpaired t-test: *p* = 0.080; n_WT_ = 7 neurons, 5 mice; n_*Fmr1*KO_ = 6, 3) **(E)** Evoked EPSC input-output slope is not significantly different for LA PNs from WT and *Fmr1*KO mice (unpaired t-test: *p* = 0.94; n_WT_ = 7 neurons, 5 mice; n_*Fmr1*KO_ = 6, 3) **(F)** Mean evoked IPSC amplitude as a function of stimulation intensity. Shading shows s.e.m. **(G)** Stimulation for half-maximal IPSC amplitude is increased for LA PNs from *Fmr1*KO mice (unpaired t-test: *p* = 0.017; n_WT_ = 7, 4; n_*Fmr1*KO_ = 7, 3). **(H)** Evoked IPSC input-output slope is not significantly different for LA PNs from WT and *Fmr1*KO mice (unpaired t-test: *p* = 0.62; n_WT_ = 7, 4; n_*Fmr1*KO_ = 7, 3). **(I)** Representative mean traces of EPSCs in paired-pulse experiments. Darker colors show 20 ms interstimulus interval, and lighter colors show 100 ms interstimulus interval **(J)** Paired-pulse ratio is increased in PNs from *Fmr1*KO mice and for shorter interstimulus interval durations (2-way repeated measures ANOVA, main effects of genotype and interstimulus interval: *p*_Genotype_ = 0.047, *p*_interstimulus interval_ = 1.56×10^−5^; nWT = 6, 4; n*Fmr1*KO = 5, 3). All summary statistics as mean ± s.e.m. *p < 0.05, ***p < 0.001.

Finally, we compared the presynaptic strength of the thalamic afferents onto LA PNs. To do this, we performed experiments where we stimulated the thalamic afferents in quick succession (20 and 100 ms interstimulus intervals) while recording EPSCs in the postsynaptic LA PN. These experiments revealed a selective increase in the paired-pulse ratio in LA PNs of *Fmr1*KO mice (Fig. 3I, J; 2-way repeated measures ANOVA, *p*_Main Effect: genotype_ = 0.047, *p*_Main Effect: interstimulus interval_ = 1.56×10^−5^). Taken together, these data indicate a specific disruption of local E/I balance caused by a reduction in FFI onto local PNs in the *Fmr1*KO LA.

### Reduced FFI enhances synaptic plasticity during early development

Local INs provide FFI onto PNs to gate LTP in BLA microcircuits (Bissière *et al.*, 2003; Tully *et al.*, 2007; Wolff *et al.*, 2014; Bazelot *et al.*, 2015), and LTP cannot be induced in PNs if local inhibition is intact (Bissière *et al.*, 2003). In light of the observed reduction in FFI onto LA PNs in *Fmr1*KO mice (Fig. 3), we hypothesized that it would be possible to induce LTP in *Fmr1*KO LA without manipulating GABAergic neurotransmission. To test this hypothesis, we recorded EPSCs in LA PNs following stimulation of the internal capsule. After a stable five minute baseline recording (stimulation frequency = 0.066 Hz), a high frequency tetanus stimulation was given to the internal capsule to induce LTP (2 trains of 100 pulses delivered at 100Hz 20 seconds apart). As expected, we found that LA PNs did not undergo LTP in slices from WT mice (Fig. 4A, B; unpaired t-tests: *p*_WT1_ = 0.61, *p*_WT2_ = 0.20, *p*_WT3_ = 0.18; see Materials & Methods for how plasticity was determined). However, when we performed these same experiments in slices from *Fmr1*KO mice, we found that all neurons underwent significant LTP following high frequency stimulation of thalamic afferents (Fig. 4A, B; *p*_*Fmr1*KO1_ = 0.0042, *p*_*Fmr1*KO2_ = 8.69×10^−4^, *p*_*Fmr1*KO3_ = 0.013). Thus, the reduced FFI onto LA PNs in *Fmr1*KO mice correlated with enhanced synaptic plasticity induction in an important circuit for classical sensory-threat conditioning.

**Figure 4.**
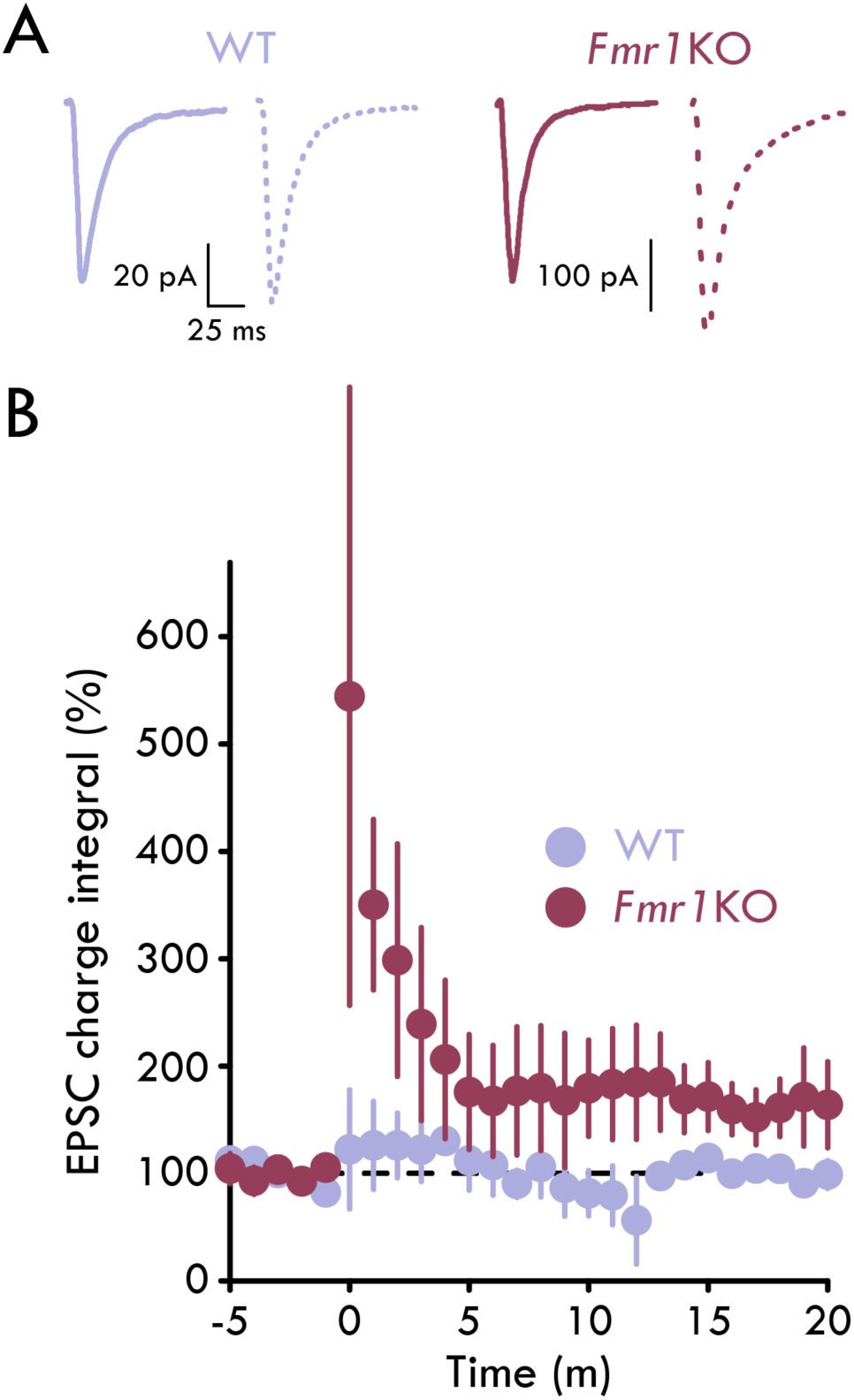
Aberrant LTP in *Fmr1*KO LA. **(A)** Representative mean EPSCs from LA PNs. Solid lines show mean EPSCs prior to LTP induction and dashed lines show EPSCs at end of experiment. **(B)** Normalized EPSC charge integral across LTP experiments. No PNs underwent significant LTP in WT LA and all PNs underwent LTP in *Fmr1*KO LA (unpaired t-tests: *p*_WT1_ = 0.61, *p*_WT2_ = 0.20, *p*_WT3_ = 0.18, *p*_*Fmr1*KO1_ = 0.0042, *p*_*Fmr1*KO2_ = 8.69×10^−4^, *p*_*Fmr1*KO3_ = 0.013; n_WT_ = 3 neurons, 2 mice; n_*Fmr1*KO_ = 3, 2). Summary statistics presented as mean ± s.e.m.

## DISCUSSION

Synaptic dysfunction is a core aspect of neurodevelopmental disorders (Chao *et al.*, 2010; Zoghbi & Bear, 2012). In the present study, we investigated circuit function in a brain region known for reduced inhibitory neurotransmission in the *Fmr1*KO mouse model of FXS. Here, we show that PNs within the LA of *Fmr1*KO mice exhibit intrinsic membrane hyperexcitability over the P21-35 developmental time point. Further, we show alterations in spontaneous and sensory afferent evoked excitatory and inhibitory neurotransmission. In particular, we demonstrate a preferential reduction in FFI onto PNs in the *Fmr1*KO LA. Finally, we find that the loss of FFI onto PNs and the increase in their excitability correlate with an enhancement of LTP at sensory afferents to the LA of *Fmr1*KO mice. Thus, we show coordinated changes in physiology, circuit function, and synaptic plasticity in a neural circuit responsible for sensory-threat learning.

### Increased excitability in LA likely contributes to adverse behavioral symptoms in FXS

Within the LA, activity-dependent excitation of PNs underlies associative threat learning (Repa *et al.*, 2001; Rosenkranz & Grace, 2002; Wolff *et al.*, 2014; Duvarci & Pare, 2014). For instance, onset of the conditioned stimulus (CS) during classical Pavlovian threat conditioning elicits strong excitation of projection PNs. Increases in CS-evoked spike activity are observed in LA PNs after training (Quirk *et al.*, 1995; Schoenbaum *et al.*, 1999). Further, threat conditioning results in enhanced excitatory synaptic transmission of the auditory thalamic afferents onto PNs of the LA (McKernan & Shinnick-Gallagher, 1997; Namburi *et al.*, 2015), and input-specific, Hebbian-like LTP underlies this synaptic strengthening (Nabavi *et al.*, 2014; Kim & Cho, 2017).

Hyperexcitable PNs and the LA have been postulated to underpin a number of the neurologic and psychiatric symptoms in FXS and ASDs, as well as other neuropsychiatric disorders, including post traumatic stress and anxiety disorders, attention-deficit/hyperactivity disorder, and substance use disorders (Posner *et al.*, 2011; Contractor *et al.*, 2015; Sharp, 2017). However, to date, few studies have focused on the amygdala of FXS patients in early life. Here, we demonstrate that PNs in the *Fmr1*KO LA exhibit marked hyperexcitability compared to WT. Specifically, LA PNs show increased maximum firing rates, a lower rheobase, and a more depolarized resting membrane potential. Thus, LA PN hyperexcitability may contribute to the clinical symptomatology of FXS. Previous studies in the hippocampus and across cortex identified similar hyperexcitable phenotypes in excitatory neurons of *Fmr1*KO mice resulting from alterations in HCN and voltage-gated Na^+^ and K^+^ ion channels (Gu *et al.*, 2007; Higgs & Spain, 2011; Gonçalves *et al.*, 2013; Zhang *et al.*, 2014; Kalmbach *et al.*, 2015; Deng & Klyachko, 2016; Routh *et al.*, 2017). To our knowledge, similar detailed ion channel studies have not been performed in the LA of *Fmr1*KO mice. Thus, whether these channelopathies are region-specific or are also present in the LA of *Fmr1*KO mice remains to be determined. Future studies will be needed to completely define the ionic mechanisms underlying intrinsic excitability in PNs in the LA (Pape & Pare, 2010; Duvarci & Pare, 2014). Importantly, these studies may reveal new therapeutic targets for the treatment of anxiety disorders in FXS, ASDs, or other neuropsychiatric disorders.

### E/I imbalance drives aberrant plasticity in LA

Globally, synaptic strength is modulated to maintain balanced excitatory and inhibitory activity within a network (Turrigiano, 1999). This synaptic scaling functions to maintain synaptic input in an activity-dependent manner (Turrigiano *et al.*, 1998; Kilman *et al.*, 2002). Our previous work has identified reductions in inhibitory synaptic efficiency and significant depletions in inhibitory function in PNs of *Fmr1*KO mice (Olmos-Serrano *et al.*, 2010; Vislay *et al.*, 2013; Martin *et al.*, 2014) during postnatal days P21-P30. Here, we extend these findings to include an enhancement of sEPSC amplitude in PNs in *Fmr1*KO LA. Taken together these data suggest a circuit phenotype of enhanced excitability.

Additionally, we identified an enhancement of the frequency of spontaneous inhibitory currents onto local PNs. Unlike our previous studies with full excitatory synapse blockade, in these studies we used recording conditions that would enable the evaluation of both excitatory and inhibitory synaptic neurotransmission within the same cell. To do this, we filled pipettes with a cesium-methanesulfonate intracellular solution which reduces resting and leak conductances and improves space-clamp. While imperfect, this method is superior to potassium-based internals for the measurement of more distal dendritic synapses that would normally be filtered (Williams & Mitchell, 2008). Thus, our synaptic recordings may have enabled better voltage control at more distal synapses, allowing us to evaluate additional sites of excitation and inhibition.

We speculate two possibilities for the role of enhancement of spontaneous, presynaptic inhibitory activity. First, it may function to compensate broadly for postsynaptic modulation of excitatory neurotransmission in a multiplicative manner (Turrigiano & Nelson, 2004) to maintain circuit homeostasis. Consistent with this, the magnitude of sEPSC amplitudes was increased in *Fmr1*KO LA PNs. Thus, increased sIPSC frequency may represent a compensatory homeostatic mechanism underlying gain control in the LA of *Fmr1*KO mice.

Alternatively, enhancement of spontaneous, presynaptic inhibitory activity may represent a homeostatic response to depleted FFI. FFI gates plasticity in the BLA circuit underlying learning of the sensory-threat associations (Bissière *et al.*, 2003; Tully *et al.*, 2007; Wolff *et al.*, 2014; Bazelot *et al.*, 2015). Importantly, disruption of this plasticity is believed to underlie the major pathophysiology of mood disorders such as anxiety and stress disorders (Duvarci & Pare, 2014). Our data reveal that at P21 evoked FFI is reduced in *Fmr1*KOs while evoked excitation is largely unaffected. Our results are in accordance with a recent study evaluating E/I conductance in the somatosensory cortex in the *Fmr1*KO mouse. In this study, a similar reduction in FFI in the L4→L2/3 feedforward circuit was observed coupled with a weaker decrease in feedforward excitation. Computational modeling using a parallel conductance model suggested that the overall net effect of this increase in E/I ratio was to maintain circuit homeostasis (Antoine *et al.*, 2019). Thus, the increase in spontaneous inhibitory neurotransmission may represent a homeostatic mechanism to compensate for a loss of FFI in the LA of the *Fmr1*KO mouse. However, other studies in the *Fmr1*KO somatosensory cortex have demonstrated similar synaptic alterations with concomitant reductions in experience-dependent plasticity (Bureau *et al.*, 2008; Harlow *et al.*, 2010). Future studies should address the idiosyncrasies of how loss of *Fmr1* affects distinct neural circuits across the brain and their corresponding behaviors.

In our previously published work, we revealed that excitatory PNs in *Fmr1* KOs display a tendency toward narrower integration windows (Martin *et al.*, 2014) that may imply decreased capacity for accurate input integration (Pouille & Scanziani, 2001; Isaacson & Scanziani, 2011) and plasticity in a circuit that is crucial for regulating fear and anxiety (Ehrlich *et al.*, 2009). Indeed, numerous studies examining the neural correlates of amygdala-based behaviors in human FXS patients and mouse models have demonstrated reductions in amygdala function. In adolescents and adults with FXS, imaging studies conducted during the presentation of fearful stimuli demonstrated attenuated amygdala activation (Hessl *et al.*, 2007, 2011; Kim *et al.*, 2014). In the mouse model of FXS, previous studies of PNs in the LA of *Fmr1*KO mice have identified impairments in LTP (extracellular field recordings) in PNs in the LA (Paradee *et al.*, 1999; Zhao *et al.*, 2005; Suvrathan *et al.*, 2010), reductions in the surface-expression of AMPA receptors (Suvrathan *et al.*, 2010), and impairments in mGluR-mediated LTP, a process which modulates LTP in LA under normal circumstances (Rodrigues *et al.*, 2002). These plasticity deficits also occur in the context of pre- and postsynaptic deficits including reductions of both the frequency and amplitude of miniature excitatory postsynaptic currents and weakened excitatory pre-synapses (Suvrathan *et al.*, 2010). Unfortunately, these plasticity studies were conducted in older animals and employed LTP induction protocols in the presence of GABA_A_ receptor blockers. Thus, we could not directly compare these data to how the fluctuations of excitation and inhibition in *Fmr1*KO mouse affects synaptic plasticity.

Since few studies have focused on emotional processing systems and how loss of the FMRP may affect circuit function and plasticity earlier in life, we evaluated LTP without GABA_A_ receptor blockers in younger animals to directly assess the inhibitory gating of synaptic plasticity in the juvenile LA of the *Fmr1*KO mouse. Surprisingly, contrary to previous reports of decreased synaptic plasticity in the LA of Fmr1 KO mice (Zhao *et al.*, 2005), we observed that reduced FFI correlates with *enhanced* synaptic plasticity in the thalamo-amygdalar circuit of juvenile *Fmr1*KO mice. Thus, increased plasticity in the circuits responsible for fear-learning may underpin the pathophysiology of anxiety disorders in FXS and ASDs in early life. However, anxiety and fear-related disorders in FXS and ASDs in later life may be mediated by other mechanisms including changes in brain-wide functional connectivity (Haberl *et al.*, 2015; Shen *et al.*, 2016) or changes in neuromodulation (Hessl *et al.*, 2002; Ghilan *et al.*, 2015).

Future studies will be necessary to evaluate whether exogenously altering E/I balance, perhaps through the enhancement of inhibition, is capable of normalizing synaptic plasticity in the LA of the *Fmr1* KO mouse Further, changes in synaptic plasticity and fear-learning throughout early development will be needed to determine if the trajectory of plastic changes seen in the juvenile BLA of *Fmr1* KO mice is pathologic or homeostatic.

## Acknowledgments

This work was supported by U.S. National Institutes of Health grants (R01 DC000566 to D.R. and R01 NS095311 to M.M.H) and U.S. National Science Foundation Graduate Research Fellowship (DGE-1553798 to E.M.G.).

## The Author contributions

Supervision, D.R. and M.M.H.; Conceptualization, E.M.G., M.N.S., D.R., and M.M.H; Investigation, E.M.G., M.N.S., C.C.R., K.K.; Software, E.M.G.; Formal Analysis, E.M.G., C.C.R.; Visualization, E.M.G., C.C.R.; Resources, D.R. and M.M.H.; Writing – Original Draft, E.M.G., M.N.S.; Writing – Review & Editing, E.M.G., M.N.S., S.M.B., D.R., and M.M.H.; Funding Acquisition, E.M.G., D.R., M.M.H.

## Declaration of interests

The authors declare no competing interests.

## Materials & Methods

### Contact for Reagent and Resource Sharing

Further information and requests for resources and reagents should be directed to and will be fulfilled by corresponding author, Molly M. Huntsman.

### Experimental Model and Subject Details

All experiments and procedures were conducted in accordance with protocols approved and reviewed by the Institutional Animal Care and Use Committee at the University of Colorado Anschutz Medical Campus, in accordance with guidelines from the National Institutes of Health. Slice electrophysiology experiments were conducted on mice aged postnatal (P) days 21-35. Experiments were conducted on both male (*Fmr1*^*-/y*^) and female (*Fmr1*^*-/-*^) mice. The following mouse lines were used in the experiments: C57Bl/6J (Jackson Laboratory #000664) and B6.129P2-Fmr1^tm1Cgr^/J (Jackson Laboratory #003025) and FVB.129P2-Pde6b^+^Tyr^c-ch^Fmr1^tm1Cgr^/J (Jackson Laboratory #004624). All mice were obtained from the Jackson Laboratory and housed in polypropylene cages with wood shavings with a modified 10/14 hour light/dark cycle. Food and water were available ad libitum.

### Acute slice preparation for electrophysiology

Mice aged P21–35 were first anesthetized with carbon dioxide (CO_2_) and decapitated. Brains were quickly removed by dissection and glued cerebellar side-down on a vibratome (Leica Biosystems, Buffalo Grove, IL, USA) stage and immersed in an ice-cold and oxygenated cutting solution (95% O_2_/5% C0_2_; in mM: sucrose, 45; glucose, 25; NaCl, 85; KCl, 2.5; NaH_2_PO_4_, 1.25; NaHCO_3_, 25; CaCl_2_, 0.5; MgCl_2_, 7; osmolality, 290-300 mOsm/kg). We prepared acute coronal slices (300 µm) containing BLA and incubated the slices in oxygenated (95% O_2_/5% CO_2_) artificial cerebrospinal fluid (ACSF; in mM: glucose, 10; NaCl, 124; KCl, 2.5; NaH_2_PO_4_, 1.25; NaHCO_3_, 25; CaCl_2_, 2; MgCl_2_, 2; osmolality 290-300 mOsm/kg) at 36°C for at least 30 minutes. All reagents were purchased from Sigma-Aldrich (St. Louis, MO, USA).

#### Electrophysiology

Slices were placed in a submerged slice chamber and perfused with ACSF heated to 32-37°C at a rate of 2 mL/min. Slices were visualized using a moving stage microscope (Scientifica: Uckfield, UK; Olympus: Tokyo, Japan) equipped with 4×(0.10 NA) and 40×(0.80 NA) objectives, differential interference contrast (DIC) optics, infrared illumination, LED illumination (CoolLED, Andover, UK), a CoolSNAP EZ camera (Photometrics, Tuscon, AZ, USA), and Micro-Manager 1.4 (Open Imaging, San Francisco, CA, USA). Whole cell patch clamp recordings were made using borosilicate glass pipettes (2.5-5.0 MW; King Precision Glass, Claremont, CA, USA) filled with an intracellular recording solution. Data was acquired with a Multiclamp 700B amplifier and were converted to a digital signal with the Digidata 1440 digitizer using pCLAMP 10.6 software (Molecular Devices, Sunnyvale, CA).

Recordings were obtained from visually identified excitatory PNs in the LA. PNs were targeted based on their large, pyramidal-like soma. Recordings were terminated if the physiology of the neuron was inconsistent with LA PNs (e.g. high membrane resistance, narrow AP halfwidth, large and fast spontaneous EPSCs).

For voltage clamp experiments, a cesium methanesulfonate (CsMe) based intracellular solution was used (in mM: CsMe, 120; HEPES, 10; EGTA, 0.5; NaCl, 8; Na-phosphocreatine, 10; QX-314, 1; MgATP, 4; Na_2_GTP, 0.4; pH to 7.3 with CsOH; osmolality adjusted to approximately 290 mOsm/kg). For all current clamp and plasticity experiments, a potassium gluconate based intracellular solution was used (in mM: potassium gluconate, 135; HEPES, 10; KCl, 20; EGTA, 0.1; MgATP, 2; Na_2_GTP, 0.3; pH to 7.3 with KOH; osmolality adjusted to approximately 295 mOsm/kg). Access resistance was monitored throughout the experiments and data were discarded if access resistance exceeded 25 MW or varied by more than 20%.

No junction potential compensation was performed. Data were sampled at 10 kHz and lowpass filtered at 4 kHz. Offline, current data were filtered using either a 3^rd^ order Savistky-Golay filter with a ±0.5 ms window or a 2 kHz lowpass butterworth filter. Mean traces were created by first aligning all events by their point of maximal rise (postsynaptic currents) and then obtaining the mean of all events.

#### Electrophysiology experimental design

*Ramped current injections* Immediately after achieving whole-cell configuration, LA neurons were recorded at rest in current clamp mode (I_hold_ = 0 pA). Following a three second baseline period, the holding current was linearly ramped from 0 pA to 400 pA over two seconds. 25 sweeps of data were collected for each neuron, and the data were used to determine the resting membrane potential, action potential threshold, and rheobase current of LA PNs.

*Square current injections* Following ramped current injections, we recorded the responses of LA neurons to a series of square hyperpolarizing and depolarizing current injections. Prior to initiation of the series of current injections, V_m_ of the LA neurons was adjusted to approximately -60 mV. Each cell was subjected to two series of 600 ms square current injections: -100 pA to +100 pA at 10 pA intervals and -200 pA to +400 pA at 25 pA intervals. The data collected in these experiments were used to determine active and passive membrane properties of the neurons.

*Spontaneous EPSC/IPSCs* Spontaneous EPSCs (V_hold_ = -70 mV) and IPSCs (V_hold_ = 0 mV) in LA PNs were recorded for 80 seconds each.

*Input-output curves* Thalamic afferents from the internal capsule were stimulated using a bipolar stimulating electrode (FHC, Inc., Bowdoin, ME, USA). We recorded evoked EPSCs (V_hold_ = -70 mV) and IPSCs (V_hold_ = 0 mV) from LA PNs in response to internal capsule stimulation. Experiments were conducted over a range of stimulation intensities (0 µA to 100 µA with a 10 µA interval).

*Paired-pulse EPSC experiments* Thalamic afferents from the internal capsule were stimulated twice at 10 Hz and 50 Hz (100 ms and 20 ms interstimulus intervals) at a 100 µA stimulus intensity. Evoked EPSCs (V_hold_ = -70 mV) from LA PNs in response to this stimulation.

*PN plasticity* For LTP experiments, we recorded PNs from WT and *Fmr1*KO mice at P21-35 (voltage-clamp configuration, V_hold_ = -80mV). AMPA mediated-excitatory postsynaptic currents (EPSCs) elicited by electrical stimulation of the internal capsule were recorded (stimulation frequency = 0.066 Hz). Following a 5 minute baseline recording, high frequency electrical stimulation (HFS; 2 trains of 100 pulses delivered at 100 Hz, 20 seconds apart) were delivered to the internal capsule. EPSCs are measured for 20-45 minutes after HFS in the same way as baseline recordings. Synaptic strength was quantified as the integrated charge of each EPSC. Change in synaptic strength was determined by normalizing the integrated charge of each EPSC recorded both before and after HFS to the average integrated charge of all baseline recordings (average normalized integrated charge of baseline = 100 %). Successful LTP induction was defined as a significant increase in normalized integrated charge during the last five minutes (minutes 16 to 20) after HFS compared to baseline (minutes -5 to -1).

#### Definitions of electrophysiological parameters

*V*_*rest*_ V_rest_ was defined as the mean V_m_ (I_hold_ = 0 pA) during a 500 ms baseline across all sweeps in the ramped injection experiments.

*AP threshold* AP threshold was defined as the voltage at which dV/dt exceeded 20 V/s. AP threshold was calculated at the first action potential of each sweep in the ramped injection experiments.

*Rheobase current* Rheobase current was defined as the mean current injected at AP threshold for the first AP across all sweeps in the ramped injection experiments.

*Membrane resistance* Membrane resistance was defined as the slope of the best fit line of the I-V plot using the -100 pA to +100 pA (10 pA steps) series of current injections. Mean voltage response to each current injection step was defined as the difference between baseline mean membrane voltage (100 ms prior to current injection) and the mean membrane voltage during the 100 ms period from 50 ms after the start of the injection to 150 ms after the start of the current injection. This 100 ms window was chosen to allow for measurement of the change in V_m_ after the membrane had charged and prior to any potential HCN channel activation. The I-V plot was constructed using all current steps below rheobase.

*Maximum firing rate* Maximum firing rate was defined as the inverse of the inter-spike interval (ISI) during the first 200 ms of the most depolarizing current injection step before attenuation of action potential (AP) firing was observed. Max FR was calculated using the -200 pA to +400 pA (25 pA steps) series of current injections.

*AP amplitude* Amplitude of the AP was defined as the voltage difference between the peak of the AP and its threshold potential (set at dV/dt = 20 V/s). AP amplitude was calculated at the rheobase sweep of the -200 pA to +400 pA (25 pA steps) series of current injections.

*AP halfwidth* AP halfwidth was defined as the time between the half-amplitude point on the upslope of the AP waveform to the half-amplitude point on the downslope of the AP waveform. AP halfwidth was calculated at the rheobase sweep of the 200 pA to +400 pA (25 pA steps) series of current injections.

*After-hyperpolarization potential (AHP) magnitude* AHP magnitude was defined as the difference between the most hyperpolarized membrane voltage of the AHP (occurring within 100 ms after AP threshold) and AP threshold. AHP magnitude and latency data were calculated at the rheobase sweep of the -200 pA to +400 pA (25 pA steps) series of current injections. ΔAHP data were calculated at the rheobase + 50 pA sweep of the -200 pA to +400 pA (25 pA steps) series of current injections.

*AHP latency* AHP latency was defined as the time from AP threshold and the peak of the AHP.

*ΔAHP* ΔAHP was defined as the difference between the first and last AHP (ΔAHP = AHP_last_ -AHP_first_).

*AP phase plot* The action potential phase plot was obtained by plotting the rate of change of the mean AP for each cell from the rheobase sweep of the -200 pA to +400 pA (25 pA steps) series of current injections as a function of the corresponding membrane voltage.

*Latency to first AP* AP latency was defined as the time from the initiation of the current injection to the peak of the first AP. AP latency was calculated at the rheobase sweep of the -200 pA to +400 pA (25 pA steps) series of current injections. *Firing rate adaptation ratio (FR adaptation).* Firing rate adaptation was defined as the ratio of the first and the average of the last two ISIs, such that Firing Rate adaptation = ISI_first_/meanISI_last two ISI_. Firing rate adaptation was calculated at the rheobase +50 pA sweep of the -200 pA to +400 pA (25 pA steps) series of current injections.

*AP broadening* AP broadening was defined as the ratio of the AP halfwidths of the first two APs (Broadening = halfwidth_second_/halfwidth_first_). AP broadening was calculated at the rheobase +50 pA sweep of the -200 pA to +400 pA (25 pA steps) series of current injections.

*AP amplitude adaptation* AP amplitude adaptation was defined as the ratio of the AP amplitude of the average of the last three APs and the first AP, such that AP Amplitude adaptation = meanAmplitude_last 3 APs_/Amplitude_first AP_. AP amplitude adaptation was calculated at the rheobase +50 pA sweep of the -200 pA to +400 pA (25 pA steps) series of current injections.

*Membrane decay τ* Membrane decay τ was determined by using a single exponential fit, f(t) = A*e*^−*t*/*τ*^ to fit the change in V_m_ induced by a -100 pA sweep in the -100 pA to +100 pA (25 pA steps) series of current injections.

*Hyperpolarization-induced sag* Hyperpolarization-induced sag was calculated using the equation, 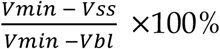, where V_min_ was defined as the most hyperpolarized membrane voltage during the current injection, V_ss_ was defined as the mean steady-state membrane voltage (last 200 ms of the current injection), and V_bl_ was defined as the mean baseline membrane voltage (100 ms prior to current injection). Hyperpolarization-induced sag was measured from the -200 pA current injection.

*Rebound spikes* Rebound spikes were defined as the number of APs in the 500 ms following the -200 pA current injection.

*sEPSC/IPSC detection and amplitude* sEPSC/IPSCs were detected by a combined template and threshold method. Briefly, a template was made by subsampling 10% of local peaks exceeding at least 6×or 7×(sEPSC or sIPSC, respectively) the median absolute deviation of a rolling baseline current (50ms prior to the peak). The template current was then truncated from its 20% rise point through the end of the decay time constant for the template current. Next, all local peaks exceeding 6×or 7×the median absolute deviation of a rolling baseline current (50ms prior to the peak) were collected. The template was then scaled to each individual putative sEPSC or sIPSC peak and each peak was assigned a normalized charge integral relative to the template. Finally, a normalized charge integral cutoff was chosen to exclude obvious noise/non-physiological events below a certain normalized charge integral. sEPSC amplitude was defined as the difference between the peak amplitude of each detected current and its corresponding baseline current. sEPSC/IPSC amplitude for each cell was defined as the median peak amplitude for that cell. sEPSC/IPSC frequency was defined as the inverse of the interevent intervals of the events. The frequency measure for each neuron was defined as the median of the sEPSC/IPSC frequencies for that cell.

*sEPSC/IPSC 20–80% risetime* 20–80% risetime was defined as the time it took an sEPSC or sIPSC to reach 80% of its peak amplitude from 20% of its peak amplitude. 20–80% risetime was calculated from the mean sEPSC/sIPSC of a given LA PN.

sEPSC/IPSC τ_Decay_. sEPSC τ_Decay_ was determined using a single exponential fit, f(t) = A*e*^−*t*/*τ*^. IPSC τ_Decay_ was defined as the weighted time-constant of IPSC decay. Briefly, a double exponential fit, f(t) = A_1_*e*^−*t*/*τ*1^+ A_2_*e*^−*t*/*τ*2^, was used to obtain the parameters to determine the weighted time-constant where τ_Weighted_ = (τ_1_A_1_ + τ_2_A_2_)/(A_1_ + A_2_). τ_Decay_ was calculated using the mean sEPSC or sIPSC trace for a cell.

*EPSC/IPSC detection and amplitude, input-output curves* To determine the evoked EPSC and IPSC amplitudes across varying stimulus intensities, we first determined the peak time relative to the 100 µA stimulation. Then, we defined EPSC or IPSC amplitude as the maximum positive or negative deflection, respectively, from the mean current response within a window of 6 standard deviations of the peak time jitter.

*Stimulation for half-maximum EPSC/IPSC amplitude* To get the half-maximum stimulation intensity and the slope of the input-output curve, we used the least squares method to fit a line to the EPSC/IPSC output relative to stimulation input. We only used input values that elicited non-zero EPSC/IPSC amplitudes to determine the best fit line. We then used this best fit line to find the stimulation intensity that was associated with 50% of the maximum EPSC/IPSC amplitude for the PN.

*Input-output curve slope* We defined input-output slope as the slope of the line created with a least squares fit of the input-output curve.

*Paired-pulse ratio* To determine paired pulse ratio, we first determined the peak amplitude of the maximum negative deflection from the mean current trace during the post-stimulus period (20ms or 100ms post stimulation) for each of the paired stimulations. We defined the paired-pulse ratio such that paired-pulse ratio = Amplitude_EPSC, second_/Amplitude_EPSC, first_.

### Quantification and Statistical Analysis

#### Statistical analyses

All data analysis was performed using custom written MATLAB code. Normality of the data were assessed using the Anderson-Darling test. For a test between two groups normal data, an unpaired t-test was used. For tests between two groups of non-normal data, a Mann-Whitney U test was used. For examination of paired pulse experiment results across genotype, a two-way repeated measures ANOVA was used. Genotype was used as the between-subjects model and interstimulus interval was used as the within-subjects model. All statistical tests were two-tailed. Unless otherwise stated, experimental numbers are reported as *n* = x, y where x is the number of neurons and y is the number of mice.

#### Data display

Data visualizations were created in MATLAB and Adobe Illustrator. Normal data are presented as the mean±s.e.m. Non-normal data are presented as the median with error bars extending along the interquartile range.

### Data and Software Availability

Data and code are available on request, and code will be made available on GitHub at https://github.com/emguthman.

## Notes

### Competing Interest Statement

The authors have declared no competing interest.

